# Data Imbalance in Drug Response Prediction – Multi-Objective Optimization Approach in Deep Learning Setting

**DOI:** 10.1101/2024.03.14.585074

**Authors:** Oleksandr Narykov, Yitan Zhu, Thomas Brettin, Yvonne A. Evrard, Alexander Partin, Fangfang Xia, Maulik Shukla, Priyanka Vasanthakumari, James H. Doroshow, Rick L. Stevens

**Affiliations:** Computing, Environment and Life Sciences, Argonne National Laboratory, Lemont, IL 60439, USA; Leidos Biomedical Research, Frederick National Laboratory for Cancer Research, Frederick, MD 21702, USA; Developmental Therapeutics Branch, National Cancer Institute, Bethesda, MD 20892, USA; Department of Computer Science, The University of Chicago, Chicago, IL 60637, USA

## Abstract

Drug response prediction (DRP) methods tackle the complex task of associating the effectiveness of small molecules with the specific genetic makeup of the patient. Anti-cancer DRP is a particularly challenging task requiring costly experiments as underlying pathogenic mechanisms are broad and associated with multiple genomic pathways. The scientific community has exerted significant efforts to generate public drug screening datasets, giving a path to various machine learning (ML) models that attempt to reason over complex data space of small compounds and biological characteristics of tumors. However, the data depth is still lacking compared to computer vision or natural language processing domains, limiting current learning capabilities. To combat this issue and increase the generalizability of the DRP models, we are exploring strategies that explicitly address the imbalance in the DRP datasets. We reframe the problem as a multi-objective optimization across multiple drugs to maximize deep learning model performance. We implement this approach by constructing Multi-Objective Optimization Regularized by Loss Entropy (MOORLE) loss function and plugging it into a Deep Learning model. We demonstrate the utility of proposed drug discovery methods and make suggestions for further potential application of the work to promote equitable outcomes in the healthcare field.

**Availability:** https://github.com/AlexandrNP/MOORLE

## 1 Introduction

Cancer is a widely spread genetic disease family with a common characteristic of uncontrolled cell growth and proliferation (Bray, et al., 2021; Cronin, et al., 2018). This set of complex genetic disorders is highly heterogeneous and notoriously difficult to combat. Artificial intelligence (AI) technologies are being incorporated into this field to facilitate our ability to treat patients. E.g., machine learning (ML) systems are usedoctors in processing radiological images and histopathological information.

Drug response prediction (DRP) is an important application of ML as it projects our estimates for the small ligand efficacy in treating cancer (Fig.1). Designing efficient DRP models can help with real-world problems of drug repurposing, personalized medicine, and virtual drug screening by reducing the number of costly wet lab experiments required to devise novel treatment protocols or develop new drugs.

**Fig. 1.**
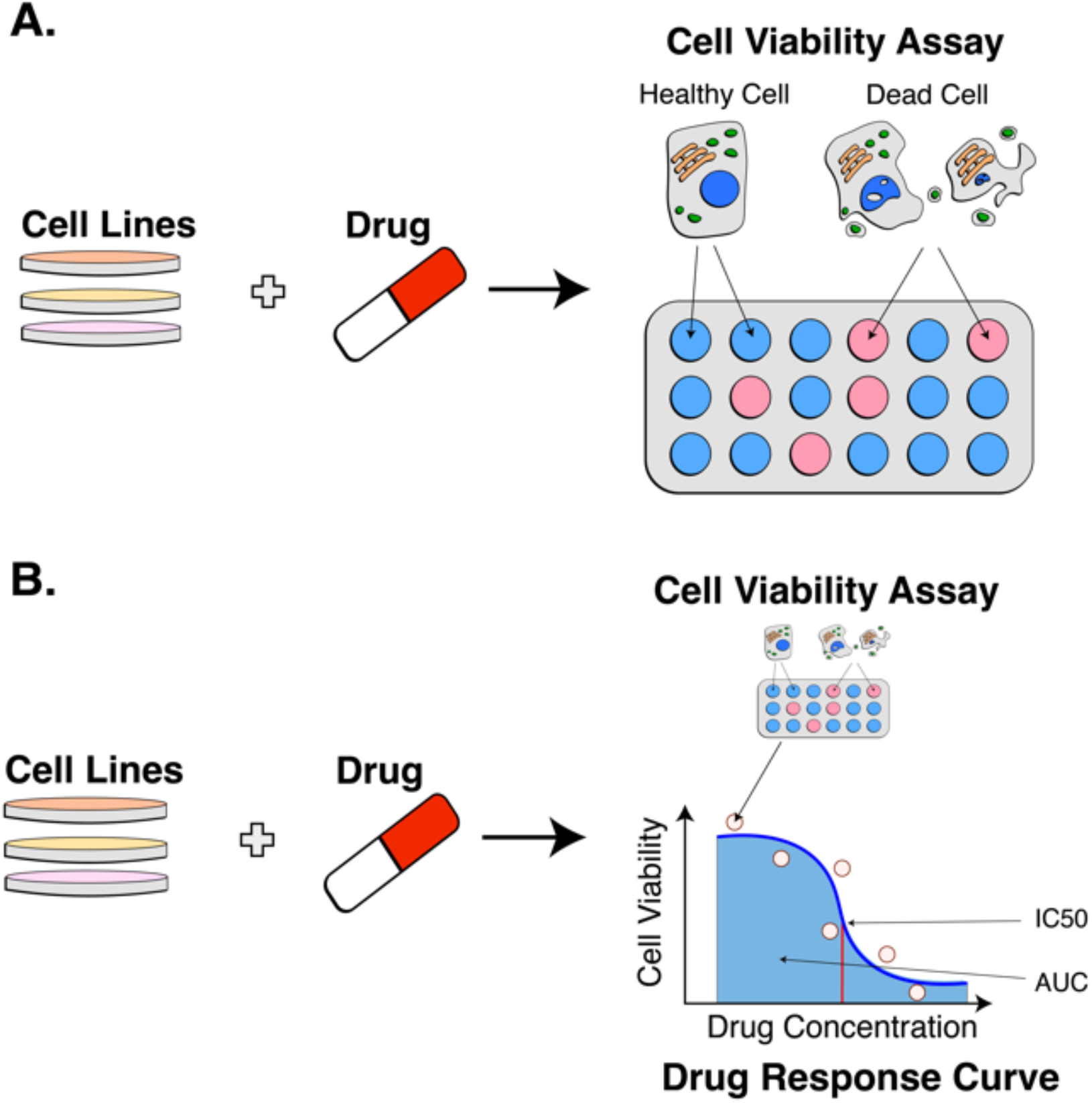
Drug response experiment. A. A single cell viability assay. A combination of membrane-permeable and membrane-impermeable fluorescent proteins can estimate the degree of tumor inhibition. B. Multiple cell viability assays are integrated into a single drug response measurement based on the hill-slope model, graphical interpretation of IC50 and AUC drug response measures (Yanagawa, et al., 1989).

However, all those settings require different approaches for assessment. Some models approach the problem in drug-specific scenarios, corresponding to personalized medicine settings, e.g., MOLI (Sharifi-Noghabi, et al., 2019). They predict drug response for a particular small molecule-based exclusively on biological information. These models cannot make inferences based on previously unseen chemical compounds. However, this setting severely limits the amount of information available for training and the strength of the model. To alleviate this issue and extend the application to different scenarios, most works approach DRP as a pair-input problem. This setting is called pan-drug DRP (Partin, et al., 2023).

The DRP field is abundant and contains multiple models based on traditional ML approaches – Random Forest (Breiman, 2001), AdaBoost (Hastie, et al., 2009), XGBoost (Lu, et al., 2021), LightGBM (Ke, et al., 2017), and Support Vector Machine (Chapelle, et al., 1999). The recent trend is the extensive usage of Deep Learning (DL) models that utilize automatic feature extraction associated with multi-layer Neural Networks (NN). One of the first approaches in this direction was described by (Menden, et al., 2013), who proposed a single-layer NN for predicting IC50. In recent years, the DRP field has had multiple models based on various architectures – convolutional neural networks (DeepIC50, DeepCDR, IGTD), graph neural networks (Graph-DRP, GraTransDRP), attention-based models (PaccMann, CADRE, DeepTTA) (Chu, et al., 2022; Jiang, et al., 2022; Joo, et al., 2019; Liu, et al., 2020; Nguyen, et al., 2021; Oskooei, et al., 2018; Tao, et al., 2020; Zhu, et al., 2021).

A pair-input setting introduces known model evaluation pitfalls (Park and Marcotte, 2012), and it is important to make appropriate train/test splits to get generalizable performance estimates. So, for a drug repurposing scenario, it is natural for the model to have prior information on both biological samples and ligands. It means that the training set may include the response of the drug in question on another cell line and the response of some other drugs on a given cell line. As long as the combination of biological data and ligands is unique, including it in the test set is appropriate. However, for virtual drug screening, ensuring that the model has no prior information on the small molecule is essential. This means that if a drug appears in a test set entry, no pairs should be involved in the training set. Otherwise, we would observe information leakage and have over-optimistic results.

Personalized medicine aims to find a treatment plan best suited for a specific patient based on their biological characteristics – genetic makeup, disease history, and style of living. DRP applications in this area aim to detect drug resistivity and, for cancer, find the most efficient drug to combat tumors specific to a given patient (Partin, et al., 2023; Zhu, et al., 2020).

Virtual screening (McGaughey, et al., 2007) setting is one of the most challenging applications for DRP models, as drug response variability between drugs is much higher than between cell line response variations (Zhu, et al., 2020). It is vital for advancing drug discovery capabilities. A significant number of works in the field focus on a one-size-fits-all optimization approach when training models. In most cases, the target is to minimize Mean Squared Error (MSE) over all pair-inputs. The performance of pan-cancer pan-drug models is commonly evaluated on a cross-validation (CV) holdout test set using performance metrics like Pearson Correlation Coefficient (PCC) and the coefficient of determination (R2). This approach assumes the ability of ML algorithms to uncover relationships between variables automatically and hides the complexity of underlying dataset structures. While multiple works address the confounding factors for the regression, most of them focus on information from different modalities, e.g., copy number variations (CNV) or mutation data (He, et al., 2022; Jia, et al., 2021; McNamee, 2005).

In this work, we are investigating the benefits of explicitly addressing complex substructures arising from p from the dataset construction with a focus on improving the virtual screening problem (Fig.2). We discuss existing approaches for learning from imbalanced data and propose an outlook on drug response prediction as a multi-objective optimization (MOO) task, attempting to maximize the prediction performance over different drugs and cancers. MOO approaches usually address problems that have multiple criteria for their evaluation. Recent work proposed its usage in contrastive learning (Moukafih, et al., 2023).

**Fig. 2.**
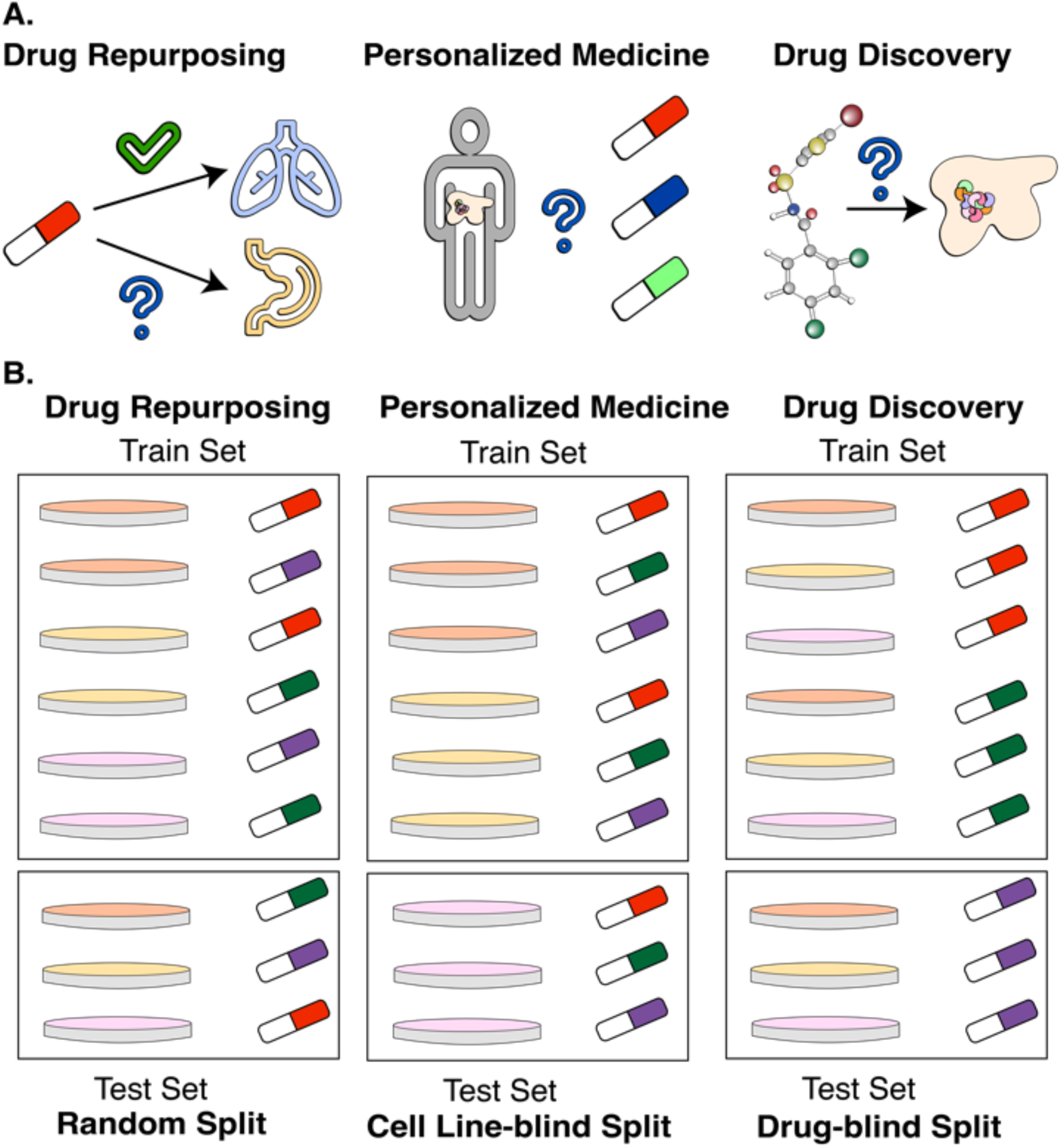
Drug response prediction application areas. A. Real-world tasks that benefit from DRP models. B. Corresponding splits of pair-input entries from drug response datasets. Each Petri dish color corresponds to a unique cell line, and each color of the drug corresponds to a unique drug.

Due to the pair-input way of constructing datasets for DRP, we can approach this problem as a hybrid formulation between classification and regression tasks. While the final goal of DRP models is to predict the continuous value of area under the drug response curve (AUC) or 50% inhibition (IC50) (Kurilov, et al., 2020), the dataset imbalance follows conducted experiments across discrete cell lines and ligand names, which can be understood as classes. Our current work focuses on drug discovery applications, corresponding to the drug-blind split of the datasets. In addition, we are also providing an assessment for the drug repurposing task to validate proposed methods, as it is one of the standard formulations of the DRP problem (Partin, et al., 2023).

## 2 Methods

### 2.1 Data

The primary data source in the DRP field is cell line experiments that measure tumor inhibition via cell viability assays. Multiple metrics characterize experimental tumor inhibition results, the most widespread being the cutoff for drug concentration that provides IC50 and AUC (Fig.1). Those are continuous metrics, so regression models are traditionally used to estimate them.

In this study, we use standard DRP cell line datasets – Cancer Therapeutics Response Portal (CTRP) (Basu, et al., 2013) and Cancer Cell Line Encyclopedia (CCLE) (Barretina, et al., 2012). CCLE dataset contains 8,950 experiments based on 474 unique cell lines and 24 drugs. Data comprises RNA-Seq gene expressions, corresponding compounds, and drug response for the combination of those two entries. The CTRP dataset does not contain gene expression data but utilizes standard commercially available cell lines and contains a much larger number of drug response experiments - 254,566. It is based on 812 cell lines and 495 ligands. Gene expression data for biological samples came from different sources, including CCLE. While dataset CCLE is mainly balanced, the variability of the experiment number is much higher for CTRP (Fig.3). Having CCLE as one of the test sets also helps us assess whether datasets with a high number of classes benefit from the proposed methodologies, regardless of the imbalance presence.

**Fig 3.**
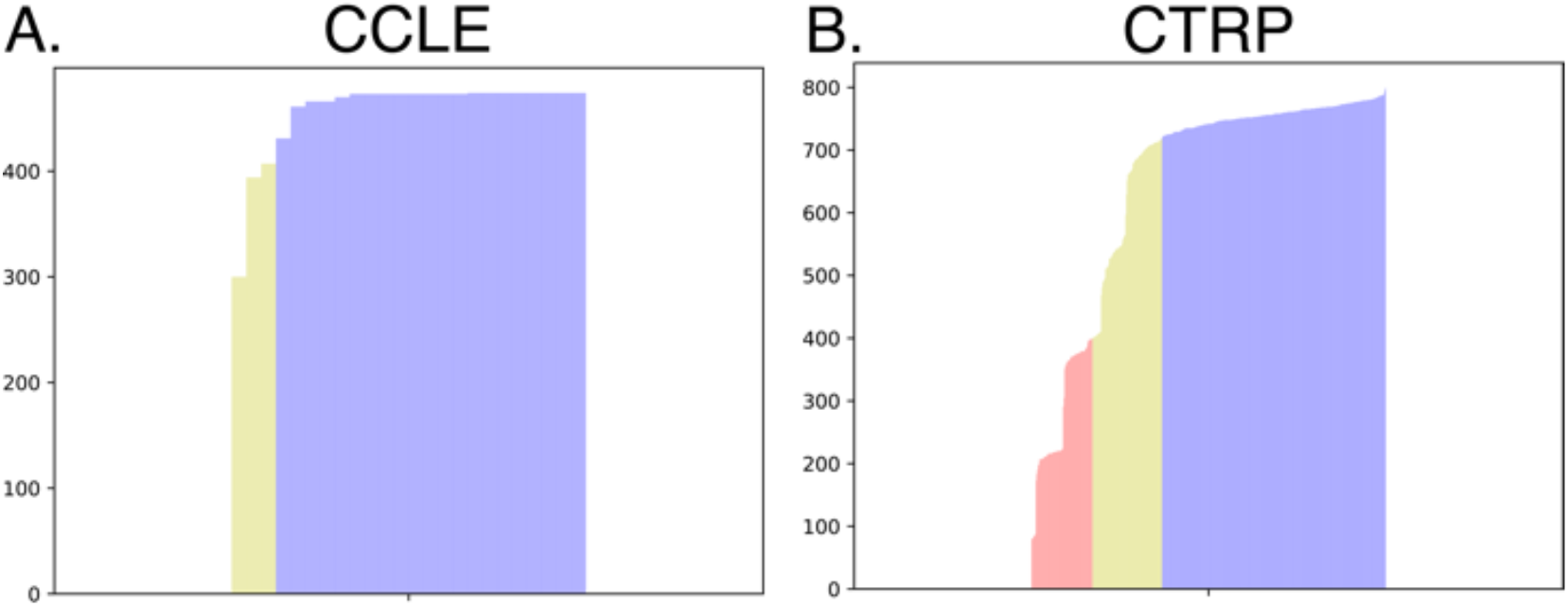
Number of experiments associated with each drug in a dataset. A. CCLE dataset. The highest number of experiments related to a single drug is 474. 12.5% of the drugs (yellow) have a number of experiments related to them, which is less than 90% of the highest number. The rest of the drugs are depicted in blue. B. CTRP dataset. The highest number of experiments associated with a single drug is 799. 17.2% of the drugs (red) have a number of experiments related to them, which is less than 50% of this highest number. 19.6% of the drugs (yellow) have a number of experiments between 50% and 90% of the highest number of experiments. The rest of the drugs are depicted in blue.

Drug-level information for both datasets comes in the form of molecular fingerprints computed via Dragon v.7.0 (Chemoinformatics) and Simplified Molecular-Input Line-Entry System (SMILES) (Weininger, 1988) entries obtained from the PubChem (Kim, et al., 2019) and the web form of Developmental Therapeutic Program (DTP) (https://dtp.cancer.gov/).

### 2.2 Learning from imbalanced data

Data imbalance is a common problem in machine learning arising due to the limited amount of available learning data (Haixiang, et al., 2017; Rezvani and Wang, 2023). This issue garnered more attention in the context of classification. Traditionally, two major approaches have been developed to handle data imbalance in the datasets: sampling (Fig.4) and cost adaptation.

**Fig. 4.**
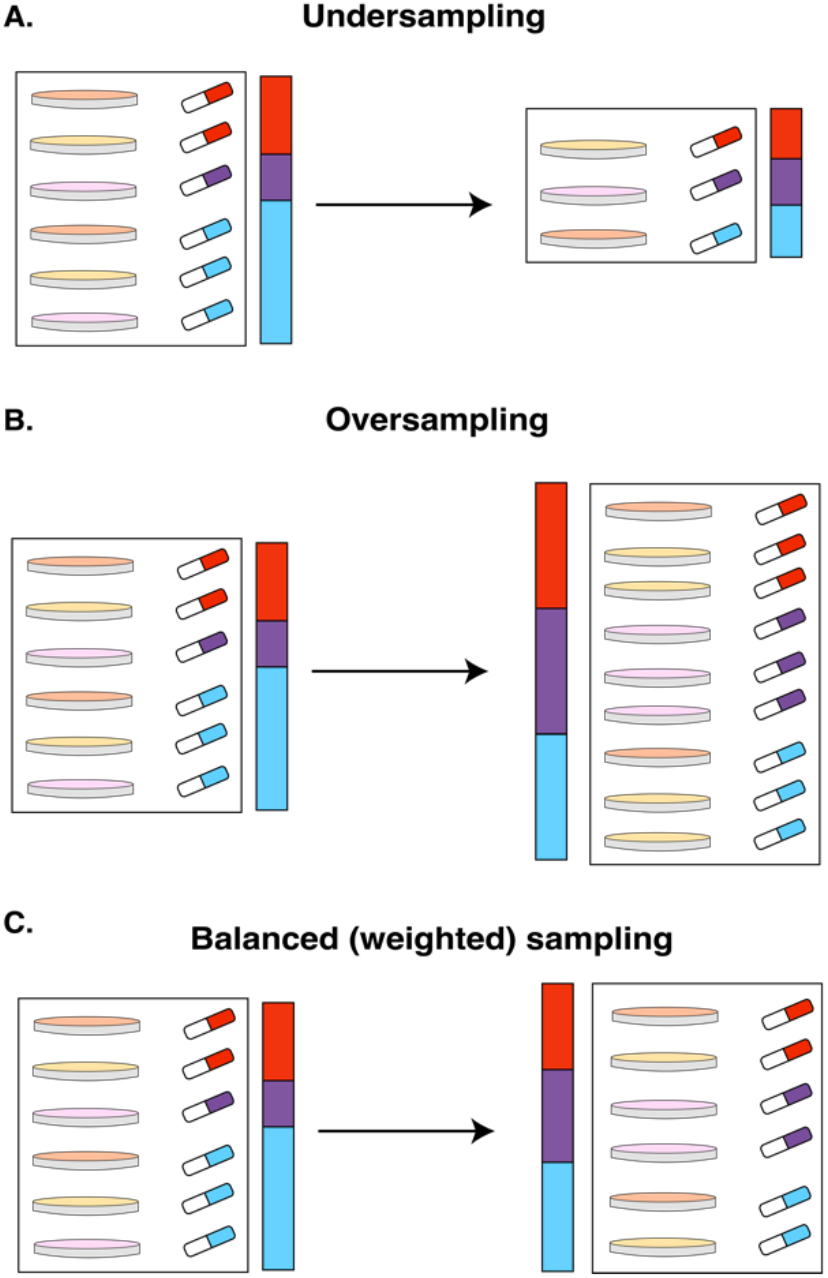
Common sampling strategies for an imbalanced dataset in the context of the DRP problem. Each rectangle represents a dataset. A Petri dish of a distinct color corresponds to the unique cell line. Drugs of different colors represent unique small molecules and compose distinct groups in data. The proportion of unique drugs is also displayed in a color bar near each dataset. In our study, we treat each drug as a class. A. Undersampling. B. Oversampling. C. Balanced, or weighted, sampling.

The first is focused on data preprocessing and includes various sampling techniques and synthetic data generation. It includes undersampling, over-sampling, a combination of these approaches (e.g., weighted sampling) (Fig.4), the SMOTE (Chawla, et al., 2002) technique, and its adaptation to the regression problem (Torgo, et al., 2013). The undersampling strategy balances data by discarding excessive data in overrepresented classes. It works well when samples of the same class are similar and additional data points from that class are not crucial for making precise predictions. It is unsuitable for the DRP problem because it leads to severe data loss. Oversampling randomly draws instances from underrepresented classes with replacement until the number of examples from each class is balanced. In this case, we have no data loss; however, the importance of data points from underrepresented classes becomes inflated, which may bias the model (Haixiang, et al., 2017). Depending on the number of classes and present imbalance, oversampling may significantly inflate the size of the dataset. Balanced (weighted) sampling can be seen as a combination of the previous two approaches. Each data point is assigned a weight inversely proportional to the number of corresponding class instances in the dataset.

The SMOTE technique is based on *k* nearest neighbors and generates synthetic examples as a weighted average between a selected point and each of its neighbors (closest data points) from the same class (Chawla, et al., 2002). The algorithm was adopted for regression; however, it heavily relies on the assumption of linearity between features and response value, as well as the convexity of the clusters formed by different classes, which are not observed in DRP problem data. It is also known that in high-dimensional space, SMOTE tends to be severely biased towards underrepresented classes (Rezvani and Wang, 2023). Because of these points, we do not further consider SMOTE in our work.

The second major approach focuses on the learning algorithm modifications. There is a large body of works for classification problems that attempts to introduce class weights (weighted variations of random forest (Breiman, 2001) and SVM (Chapelle, et al., 1999), modify the loss function, (particleswarm optimization network (Cao, et al., 2013), zSVM (Abarzadeh, et al., 2016), or refine boosting approaches (AdaC1-AdaC3 (Raghuwanshi and Shukla, 2019), RareBoost (Joshi, et al., 2001), BABoost(Song, et al., 2009). For regression, probability-based methods such as reframing were introduced. This approach focuses on adapting to estimated outputs depending on the context (Hern’ ndez-Orallo, 2014).

### 2.3 Drug Response Prediction as Multiple Objective Optimization

As we discussed earlier, the most common model evaluation is based on integral performance, e.g., *R*^2^, PCC, concordance index, etc. In this paragraph, we are using *R*^2^ as an example of performance measurement, and we refer to the standard evaluation of the entire hold-out portion of the dataset as 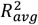 . This measure compares estimates of the residual sum of squares produced by the model

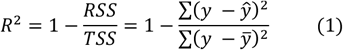

where *y* is the ground truth value, ŷ is the model prediction, and 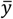 is the expected value of the response variable in the test dataset. Maximizing *R*^2^ is a common target for DRP models, and can be considered a single-objective optimization problem. However, directly using *R*^2^ for training the ML algorithms is not a common approach, as the coefficient of determination is not a convex function. It is more feasible to disregard the total sum of squares and solve the optimization problem directly for the residual sum of squares. It results in a common mean squared error (MSE) loss function:

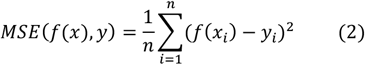

where x is the set of features, *n* – total number of datapoints, *f* – prediction model, *y* - ground truth value.

This formulation is suitable for drug repurposing task, as it considers each unique combination of biological sample and ligand a unique standalone data sample. However, when we discuss the virtual drug screening application, we are interested in the model’s ability to reason over individual small molecules. It means that the performance of each drug in the dataset can be considered a standalone optimization problem. Let’s consider that for each drug we are attempting to maximize 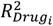 . We can construct a data space where each coefficient of determination corresponding to *i*-th drug forms an orthonormal basis. Then each individual machine learning model can be uniquely described based on the performance it achieves for the corresponding small molecule (Fig.5A). E.g., in two dimensional case with two drugs *Drug*_1_ and *Drug*_2_ vector 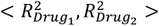 defines the coordinates of the corresponding machine learning model. On top of individual decomposition into performance evaluations 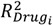 we can also associate an integral performance metric with each machine learning model. It can be either *R*^2^ over all datapoint or an average of individual performances 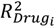 . In this work, we choose the latter because this measure is less sensitive to the number of experiments associated with each separate drug.

**Fig. 5.**
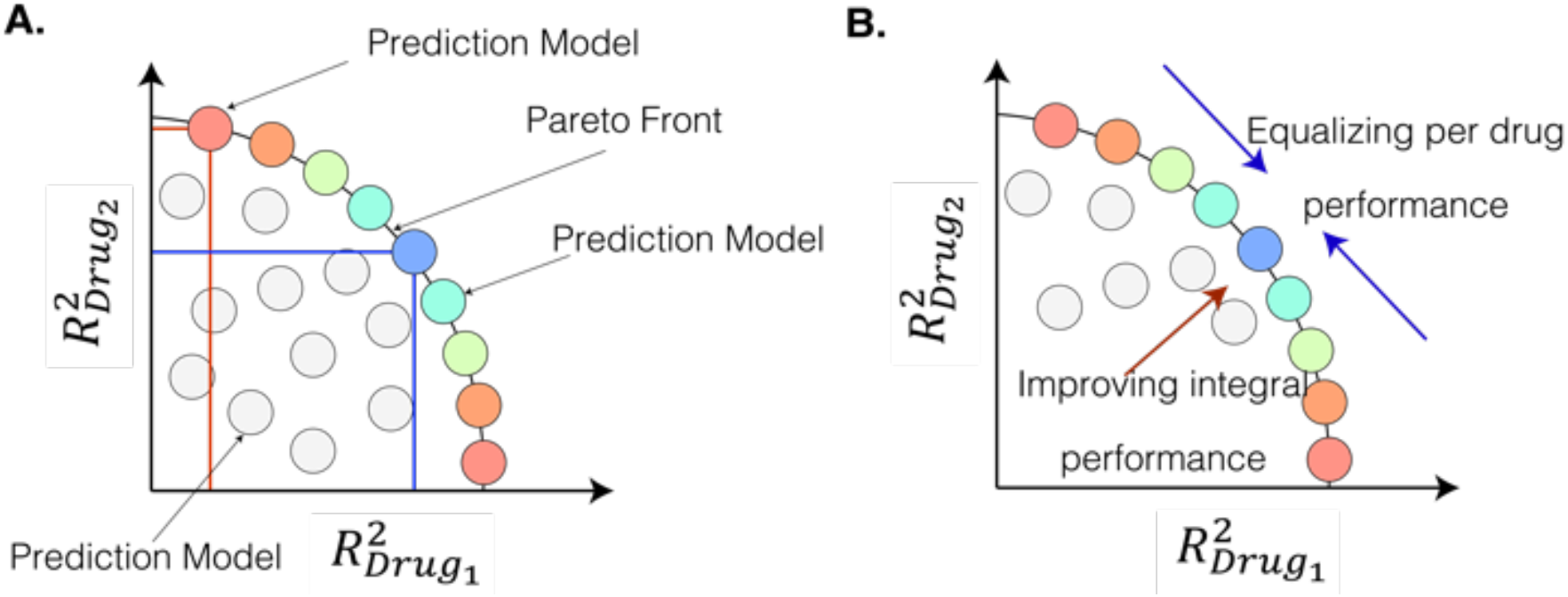
Pareto front of ML models in the space composed of individual drug performance metrics. Colored nodes indicate ML models belonging to the Pareto front, i.e., having integral performance close to the maximum known value. Grey points represent ML models that have worse integral performance.

We can see (Fig.5A) that under these assumptions, multiple data points may correspond to the machine learning models with the same integral performance but different tradeoffs between individual drugs. This is a subset of the class-composed Pareto front that is defined as all possible models with extreme performance (Teich, 2001). Now, an important question is which ML model from the Pareto front is preferable. We hypothesize that selecting models closer to the center of this set (cyan and green points from Fig.5) will result in better generalization - a capability of predicting responses to new drugs not included in the training set - as the trained models are not biased towards some drugs and thus provide a better generalizability between drugs. The reason is that a better association behind unique chemical characteristics of the small compounds and biological samples features. Preliminary findings were discussed in (Narykov, et al., 2023). Fig.5B provides a graphical visualization of our objective – to maximize the integral performance of the model and to balance individual drug scores.

As in the case with the regular *R*^2^, the current formulation does not fit to be directly used for ML algorithms training. To realize this strategy, we can define a loss function based on the MSE of individual drugs and an entropy-like regularization component. We will define loss for individual drugs as

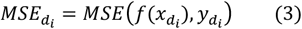

where x is the set of features, *d*_*i*_ is the *i*-th drug, *f*(. ) is the prediction model that produces AUC, *y* is the ground truth values. Then, we will calculate normalized losses by applying the softmax function to individual scores and put them in the set *P*:

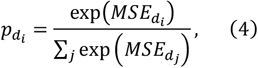

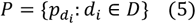

where *D* is the set of all drugs and 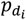 is normalized loss for drug *d*_*i*_. As we can see, set *P* can be treated as a probability distribution. As we want to incentivize equal loss for individual drugs, we can use a regularization based on the entropy function:

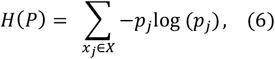

However, the function *H*(*P*) is maximized when our desired property is achieved, and is concave. So, in order to adopt it for a loss function we will use it in form ln|*D*| − *H*(*P*), where ln|*D*| is the maximum value that discrete entropy can take for the distribution with |*D*| entries. This transformation minimizes loss when drug-specific losses are equal and results in a convex function.

We will define the loss function over the set of features *x*, response values *y*, and classes *D* as

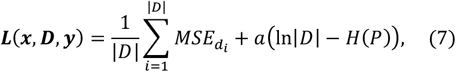

where *a* is a regularization coefficient, *H*(X) is the entropy of a distribution. The averaged sum of drug-specific losses maximizes integral performance, while the entropy-based regularization component promotes evening-out loss across drugs. We call this construction Multi-Objective Optimization Regularized by Loss Entropy (MOORLE) loss function.

Each individual part of the equation (7) is convex. It allows us to state that ***L***(***x, D, y***) function is also convex, as it consists of a linear combination of convex functions. That is not a required but desired property for loss function. It results in a versatile model-agnostic loss function that can be utilized both in classical ML models and deep learning settings.

### 2.4 Mixed Sampling Approach

The practical consideration regarding loss function modification proposed in 2.3 is the modern approach for NN training. Instead of updating gradients for the entire dataset in the gradient descent (GD) algorithm, continuous updates of model weights are made based on mini-batches. This approach allows to train models significantly faster and is a main feature of stochastic gradient descent (SGD) or its improvements, e.g., ADAM (Kingma and Ba, 2014).

When we use sequential shuffled sampling that draws each element from the dataset once in random order, and is de facto a standard for deep learning frameworks like PyTorch and Keras, there is a high possibility that underrepresented drugs would have only a small influence on the objective function. Because of these considerations, we explored different sampling strategies described in 2.2. Undersampling, as expected, led to unsatisfactory performance for both CCLE and CTRP datasets, as most of the data was discarded. Oversampling was suitable for smaller-sized CCLE, but in the case of CTRP, this approach inflated the training dataset 20 times its original size and was not computationally feasible. Weighted sampling allowed us to control the size of the training dataset manually. However, maintaining the dataset size two to four times the original training set resulted in data loss and performance deterioration.

This led us to develop a hybrid sampling strategy. Based on the common training dataset, we are deriving two sets of batches – one is the balanced batch based on weighted sampling, and another is a batch composed via sequential shuffle split (Fig.6). This mixed strategy inflates the size of the original training set twice and allows us to present model all samples from training dataset while ensuring regular update of cost function based on all present groups (classes) in the dataset. In practice, performing weighted sampling for each epoch significantly increases training time. To offset the cost of performing it, we cache obtained batches and store them for 10 epochs (including current), shuffling batch order before each new epoch. Using this heuristic, we reduce the number of times we have to perform weighted sampling.

**Fig. 6.**
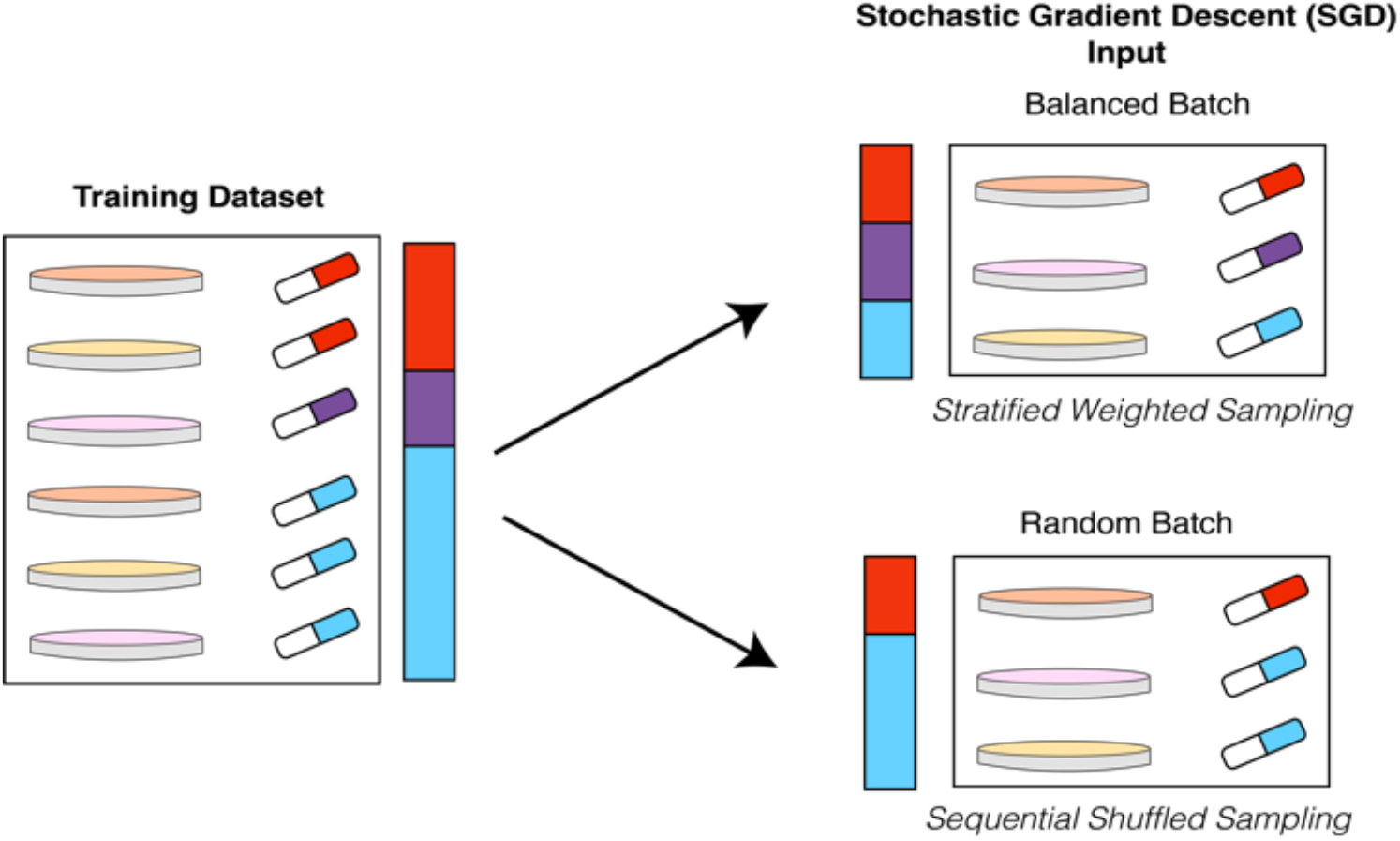
Mixed sampling scheme for stochastic gradient descent-based algorithms.

### 2.5 Machine Learning Algorithms

The approach described in section 2.3 is model-agnostic and can be potentially adopted for any ML algorithm; due to the latest trends in the field, we are focusing on its adaptation for Deep Learning settings. We incorporate loss function (7) into the recent state-of-the-art model DeepTTA (Jiang, et al., 2022).

DeepTTA consists of three main components. One is an attention-based SMILES encoder subnetwork, and the other is a fully connected neural network (FCNN) (Zhu, et al., 2021). The last part of the network concatenates encodings for drugs and biological samples and performs regression.

## 3 Results

### 3.1 Experimental setup

We analyze the effect of adopting a mixed sampling approach and entropy-regularized loss function in DeepTTA models under random split and drug-blind split CV model evaluation strategies (Fig.2B). In a drug-blind setting, a ligand can not appear in the training and testing sets simultaneously. It ensures that no information about particular ligand is present in the test set. Each model run is evaluated by 10-fold cross-validation with fixed split that is shared among the studies for the same dataset and the same split type.

We perform an ablation study to investigate the influence of sampling strategy and loss function on model performance under various conditions. We consider two sampling strategies – standard sequential and mixed sampling-proposed in Section 2.4. We also consider two loss functions – widely used MSE and multi-objective loss function with entropy regularization proposed in Section 2.3. It results in four possible combinations of the influencing factors for each model run.

To estimate the statistical significance of the effect that proposed strategies have on the drugs we apply two-way repeated measurements ANOVA (Potvin and Schutz, 2000) algorithm from *pengouin* Python package. For random split and drug-blind settings, results for each CV iteration are considered repeated measurement; for drug-averaged drug-blind settings, each individual drug plays this role. Greenhouse-Geisser corrected p-values (Abdi, 2010) are reported in the findings. Data from Fig. 7 is available in Supplementary Table 1.

**Fig. 7.**
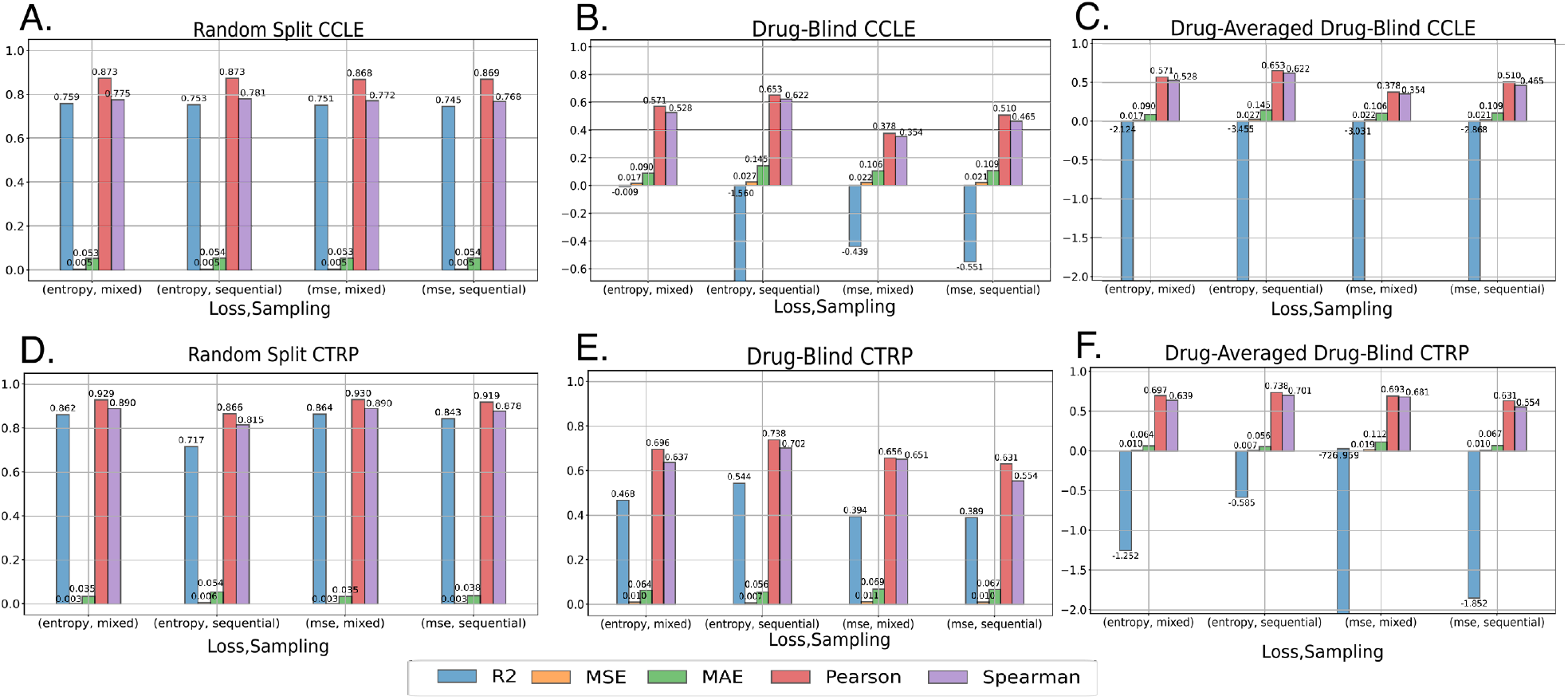
Ablation study on CCLE and CTRP datasets. Performance metrics coefficient of determination (R^2^), mean squared error (MSE), mean absolute error (MAE), Pearson correlation coefficient (Pearson), and Spearman correlation coefficient (Spearman) are recorded for each combination of factors – sampling strategy and loss function. Sampling strategies consist of sequential random sampling (denoted ‘sequential’ in the figure) and hybrid strategy introduced in 2.4 (‘mixed’ in the figure). Loss functions are represented by MSE (‘mse’ in the figure) and MOORLE - multi-objective loss function regularized by loss entropy (‘entropy’ in the figure). A. Random split evaluation strategy, CCLE dataset. B. Drug-blind split evaluation strategy, CCLE dataset. C. Drugwise evaluation under drug-blind split, CCLE dataset. D. Random split evaluation strategy, CTRP dataset. E. Drug-blind split setting, CTRP dataset. F. Drugwise evaluation under drug-blind split, CTRP dataset.

### 3.2 Random split evaluation (drug repurposing)

The random split was mainly introduced as a baseline setting to observe model behavior under the standard for the field experiment setup. We expected that the proposed change in the objective would not significantly influence this scenario, as the model has abundant information on the related ligands, unless we encounter a high imbalance of drug representations in batches. Indeed, as we can see, for CCLE (Fig.7A), with R^2^ varying from 0.745 to 0.759 and MSE staying around 0.005, while two-way repeated measurements ANOVA did not detect statistically significant effects of loss function or sampling strategy on the performance. The situation with CTRP (Fig.7B) is very similar, except for the combination of multi-objective function and sequential sampling strategy. We hypothesize that due to the completely random proportion of classes (drugs) in training batches, the model has difficulties to learn the current objective (drug repurposing)

### 3.3 Drug-blind evaluation (virtual screening, bulk assessment)

As described in Fig.2, drug-blind evaluation corresponds to the virtual screening problem, where we assess the ligand’s performance previously not seen by the model. In this scenario, performance metrics are calculated for the entire hold-out portion of the cross-validation set.

As drug-blind setup is much more challenging for ML model, we see a sharp drop in the performance. For the CCLE dataset, the best-performing combination is MOORLE loss function with mixed sampling with MAE = 0.09, while the rest of the values are 0.09, 0.11, 0.11 for MOORLE with sequential sampling, MSE with mixed sampling, and MSE with sequential sampling for CTRP dataset. We see a combination of sequential sampling and MOORLE loss function having a slight edge over the other variations with MAE = 0.056 against 0.064, 0.112, 0.067 for MOORLE with mixed sampling, MSE with mixed sampling, and MSE with sequential sampling for CTRP dataset.

For CCLE dataset, two-way repeated measurements ANOVA test highlight sampling strategy as a statistically significant performance factor (bulletsp − vaIue = 2.88 · 10^−2^) for R^2^, as well as interaction of sample strategy and loss function (p − vaIue = 6.93 · 10^−23^). For MSE, the same combination of two factors mentioned above is statistically significant (p − vaIue = 3.18 · 10^−22^). For CTRP, both sampling and loss function were not a statistically significant factors for all performance metrics.

### 3.4 Drugwise scoring under drug-blind split (virtual screening, drug-specific assessment)

Most works that discuss drug-blind evaluation perform bulk assessment, as described in 3.3. However, to be completely thorough with our assessment, we first attempt to calculate the corresponding metric for each drug individually and then average the results. As we can see from Fig.7F and Fig.7E, the only measure significantly impacted by this procedure change is R^2^. However, even when the mean values of the majority of the performance metrics stay the same, taking a look at the problem from drug-by-drug perspective allows us to better reason over the influence that changes in ML model have on the performance. As MAE score remains very close to the previous scenario, we are making comparisons based on R^2^ in this section.

For the CCLE dataset, the best-performing combination is MOORLE loss function with mixed sampling with R^2^= -2.12, while the rest of the values are -3.46, -3.03, -2.87 for MOORLE with sequential sampling, MSE with mixed sampling, and MSE with sequential sampling for CTRP dataset. We see a combination of sequential sampling and MOORLE loss function having a slight edge over the other variations with R^2^= -0.59 against -1.25, -726.96, -1.85 for MOORLE with mixed sampling, MSE with mixed sampling, and MSE with sequential sampling for CTRP dataset.

For the drug-averaged evaluation, two-way repeated measurements ANOVA test corroborated the statistically significant effect of adding both a new sampling strategy and the loss function on MSE value (p − vaIue = 1.56 · 10^−23^), and the outstanding impact of loss function on Pearson correlation coefficient (p − vaIue = 9.16 · 10^−24^) in CCLE dataset case (Fig.7C). For CTRP, the most influential factor on MSE value was sampling (p − vaIue = 3.73 · 10^−25^), with loss function also playing a significant role (p − vaIue = 4.1 · 10^−24^) (Fig.7F). As R^2^ metric is unbound on the left, it exhibits great variability across individual drugs, which we can see in the examples of top-performing and worst-performing drugs from the Table 1.

**Table 1.**
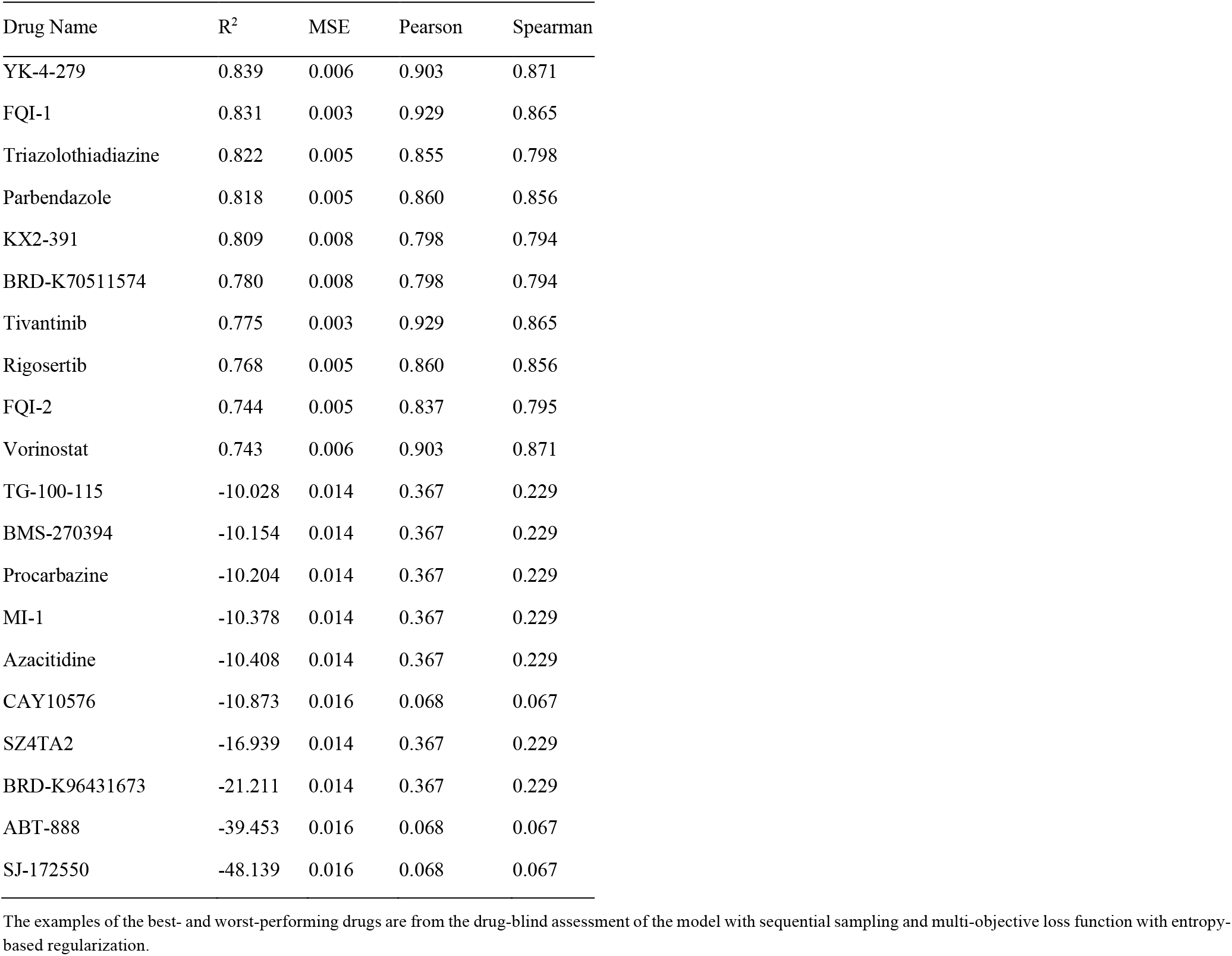
Ten drugs with the best and ten with the worst R^2^ score from the CTRP dataset.

| Drug Name | $R^2$ | MSE | Pearson | Spearman |
| --- | --- | --- | --- | --- |
| YK-4-279 | 0.839 | 0.006 | 0.903 | 0.871 |
| FQI-1 | 0.831 | 0.003 | 0.929 | 0.865 |
| Triazolothiadiazine | 0.822 | 0.005 | 0.855 | 0.798 |
| Parbendazole | 0.818 | 0.005 | 0.860 | 0.856 |
| KX2-391 | 0.809 | 0.008 | 0.798 | 0.794 |
| BRD-K70511574 | 0.780 | 0.008 | 0.798 | 0.794 |
| Tivantinib | 0.775 | 0.003 | 0.929 | 0.865 |
| Rigosertib | 0.768 | 0.005 | 0.860 | 0.856 |
| FQI-2 | 0.744 | 0.005 | 0.837 | 0.795 |
| Vorinostat | 0.743 | 0.006 | 0.903 | 0.871 |
| TG-100-115 | -10.028 | 0.014 | 0.367 | 0.229 |
| BMS-270394 | -10.154 | 0.014 | 0.367 | 0.229 |
| Procabazine | -10.204 | 0.014 | 0.367 | 0.229 |
| MI-1 | -10.378 | 0.014 | 0.367 | 0.229 |
| Azacitidine | -10.408 | 0.014 | 0.367 | 0.229 |
| CAY10576 | -10.873 | 0.016 | 0.068 | 0.067 |
| SZ4TA2 | -16.939 | 0.014 | 0.367 | 0.229 |
| BRD-K96431673 | -21.211 | 0.014 | 0.367 | 0.229 |
| ABT-888 | -39.453 | 0.016 | 0.068 | 0.067 |
| SJ-172550 | -48.139 | 0.016 | 0.068 | 0.067 |

The examples of the best- and worst-performing drugs are from the drug-blind assessment of the model with sequential sampling and multi-objective loss function with entropy-based regularization.

## 4 Discussion

Current drug response prediction approaches implicitly rely on the ability of deep learning algorithms to find a true relationship between features and response values while simultaneously correcting for biases present in the dataset. However, as the amount of available data for this biomedical problem is limited, we investigated potential ways to improve predictive value by explicitly addressing the data imbalance problem.

The proposed multi-objective loss function with entropy regularization is model-agnostic and can be utilized both with classical ML algorithms and in Deep Learning. Classes, or domains ***D***, in the MOORLE loss function ***L***(***x, D, y***) are interchangeable and can guide model to be aware of other biases encoded in data, e.g., gender, race, or age that regularly appear in biomedical datasets due to present sources of bias in our society. The proposed approach can be used to promote equitable outcomes in healthcare models.

One of the drawbacks of the proposed methodology is the dependency on the regularization coefficient. It should be derived in the inner cross-validation loop to achieve the best possible performance. However, nested cross-validation for the proposed datasets is exceptionally computationally demanding, increasing the runtime by the order of magnitude. Another limitation of this work is performing an ablation study on a single model. Our future direction is to perform a large-scale analysis of the community models with multiple loss functions and hyperparameter optimization.

Surprisingly, the mixed sampling strategy positively impacted the small, better-balanced CCLE dataset and was mostly detrimental to CTRP. It is possible that because of the large number of drugs in the latter, each balanced batch did not contain enough representative samples for the corresponding drug (class). Further adjustments in controlling the number of classes sampled in a single batch are needed. At the same time, the multi-objective loss function with entropy regularization was the primary influence for the CTRP dataset.

