## Supplementary Table 1 for "Data Imbalance in Drug Response Prediction – Multi-Objective Optimization Approach in Deep Learning Setting"

**Supplementary Table 1. Results of the ablation study.** This table contains numerical values for the data presented in Fig. 6 of the main manuscript.

| Evaluation Strategy | Dataset | Sampling | Loss | R2 | MSE | MAE | Pearson | Spearman |
|---|---|---|---|---|---|---|---|---|
| Random Split | CCLE | mixed | entropy | 0.759 | 0.005 | 0.053 | 0.873 | 0.775 |
| | | mixed | mse | 0.751 | 0.005 | 0.053 | 0.868 | 0.772 |
| | | sequential | entropy | 0.753 | 0.005 | 0.054 | 0.873 | 0.781 |
| | | sequential | mse | 0.745 | 0.005 | 0.054 | 0.869 | 0.768 |
| | CTRP | mixed | entropy | 0.862 | 0.003 | 0.035 | 0.929 | 0.890 |
| | | mixed | mse | 0.864 | 0.003 | 0.035 | 0.930 | 0.890 |
| | | sequential | entropy | 0.717 | 0.006 | 0.054 | 0.866 | 0.815 |
| | | sequential | mse | 0.843 | 0.003 | 0.038 | 0.919 | 0.878 |
| Drug Blind | CCLE | mixed | entropy | -0.009 | 0.017 | 0.090 | 0.571 | 0.528 |
| | | mixed | mse | -0.439 | 0.022 | 0.106 | 0.378 | 0.354 |
| | | sequential | entropy | -1.560 | 0.027 | 0.145 | 0.653 | 0.622 |
| | | sequential | mse | -0.551 | 0.021 | 0.109 | 0.510 | 0.465 |
| | CTRP | mixed | entropy | 0.468 | 0.010 | 0.064 | 0.696 | 0.637 |
| | | mixed | mse | 0.394 | 0.011 | 0.069 | 0.656 | 0.651 |
| | | sequential | entropy | 0.544 | 0.007 | 0.056 | 0.738 | 0.702 |
| | | sequential | mse | 0.389 | 0.010 | 0.067 | 0.631 | 0.554 |
| Drug Blind Drugwise Results | CCLE | mixed | entropy | -2.124 | 0.017 | 0.090 | 0.571 | 0.528 |
| | | mixed | mse | -3.031 | 0.022 | 0.106 | 0.378 | 0.354 |
| | | sequential | entropy | -3.455 | 0.027 | 0.145 | 0.653 | 0.622 |
| | | sequential | mse | -2.868 | 0.021 | 0.109 | 0.510 | 0.465 |
| | CTRP | mixed | entropy | -1.252 | 0.010 | 0.064 | 0.697 | 0.639 |
| | | mixed | mse | -726.95 | 0.019 | 0.112 | 0.693 | 0.681 |
| | | sequential | entropy | -0.585 | 0.007 | 0.056 | 0.738 | 0.701 |
| | | sequential | mse | -1.852 | 0.010 | 0.067 | 0.631 | 0.554 |

**Supplementary Table 2**

**Supplementary Table 2. ML algorithm parameters**

| |
|---|
| use_lincs = False |
| target_id = 'AUC' |
| transformer_emb_size_drug = 128 |
| dropout = 0.2 |
| transformer_n_layer_drug = 8 |
| transformer_intermediate_size_drug = 512 |
| transformer_num_attention_heads_drug = 8 |
| transformer_attention_probs_dropout = 0.1 |
| transformer_hidden_dropout_rate = 0.1 |
| learning_rate = 1e-4 |
| optimizer='adam' |
| batch_size = 256 |
| epochs = 100 |
| input_dim_drug_classifier = 128 |
| input_dim_gene_classifier = 256 |
| input_dim_binding_classifier = 64 |
